# HIV Env trimers elicit NHP apex cross-neutralizing antibodies mimicking human bNAbs

**DOI:** 10.1101/2025.09.19.677470

**Authors:** Javier Guenaga, Monika Adori, Shridhar Bale, Swastik Phulera, Ioannis Zygouras, Fabian-Alexander Schleich, Xaquin Castro Dopico, Sashank Agrawal, Miyo Ota, Richard Wilson, Jocelyn Cluff, Tamar Dzvelaia, Marco Mandolesi, Agnes Walsh, Mariane B. Melo, Laurent Verkoczy, Darrell J. Irvine, Martin Corcoran, Ian A. Wilson, Diane Carnathan, Guido Silvestri, Andrew B. Ward, Gabriel Ozorowski, Gunilla B. Karlsson Hedestam, Richard T. Wyatt

**Affiliations:** Department of Immunology and Microbiology, The Scripps Research Institute, La Jolla, CA, USA; Department of Microbiology, Tumor and Cell Biology, Karolinska Institutet, SE-171 77 Stockholm, Sweden; Department of Integrative Structural and Computational Biology, The Scripps Research Institute, La Jolla, CA, USA; Vaccine Research Center, National Institute of Allergy and Infectious Diseases, National Institutes of Health, Bethesda, MD, USA; Division of Microbiology and Immunology, Emory National Primate Research Center, Emory University, Atlanta, GA, USA; Applied Biomedical Science Institute, 11011 Via Frontera, San Diego, CA 92127; Howard Hughes Medical Institute, 4000 Jones Bridge Rd., Chevy Chase, MD, 20815, USA

**Keywords:** Vaccination, HIV-1 Env apex-targeting, monoclonal antibodies, cross-neutralization, rhesus macaques

## Abstract

As a chronically replicating virus, HIV has evolved extreme sequence variability and effective shielding of functionally constrained spike protein determinants by host-derived glycans. Broadly neutralizing antibodies (bNAbs), though rare, can be isolated from people living with HIV, revealing conserved Env sites as key targets for vaccine development. One such target is the apex of the envelope glycoprotein (Env) spike. Here, we identified a vaccination strategy using heterologous HIV Env trimers covalently coupled to liposomes for multivalent display that resulted in the elicitation of cross-neutralizing HIV serum antibody responses in all immunized non-human primates (NHPs). Critically, we isolated a set of monoclonal antibodies (mAbs) that cross-neutralized multiple divergent HIV clinical isolates. High-resolution cryoEM structural analysis of mAbs from three different NHPs demonstrated that they targeted the Env trimer apex in a manner remarkably similar to that of the human infection-elicited, apex-directed bNAb PG9, representing a substantial advance in HIV vaccine development.

## Introduction

The development of an effective prophylactic HIV vaccine capable of reducing the spread of the virus remains a high priority public health goal. However, chronic HIV replication in infected hosts drives immune evasion, ^1–3^ making vaccine development extremely challenging. The sole neutralizing determinant on the surface of the virus is the trimeric envelope glycoprotein (Env) spike (Figure 1A), which has evolved to evade host antibody responses by incorporating host-derived N-glycans and altering variable domains during replication.^4,5^ Broadly neutralizing antibodies (bNAbs), which arise infrequently in people living with HIV-1 (PLWH), can penetrate and, in some cases, co-recognize components of the glycan shield and underlying protein structures, thereby revealing sites of vulnerability. These include the Env trimer apex (Figure 1A), the primary CD4 receptor binding site, V3 glycan site, and functionally constrained regions of the HIV spike proximal to the viral membrane.^6,7^

**Figure 1.**
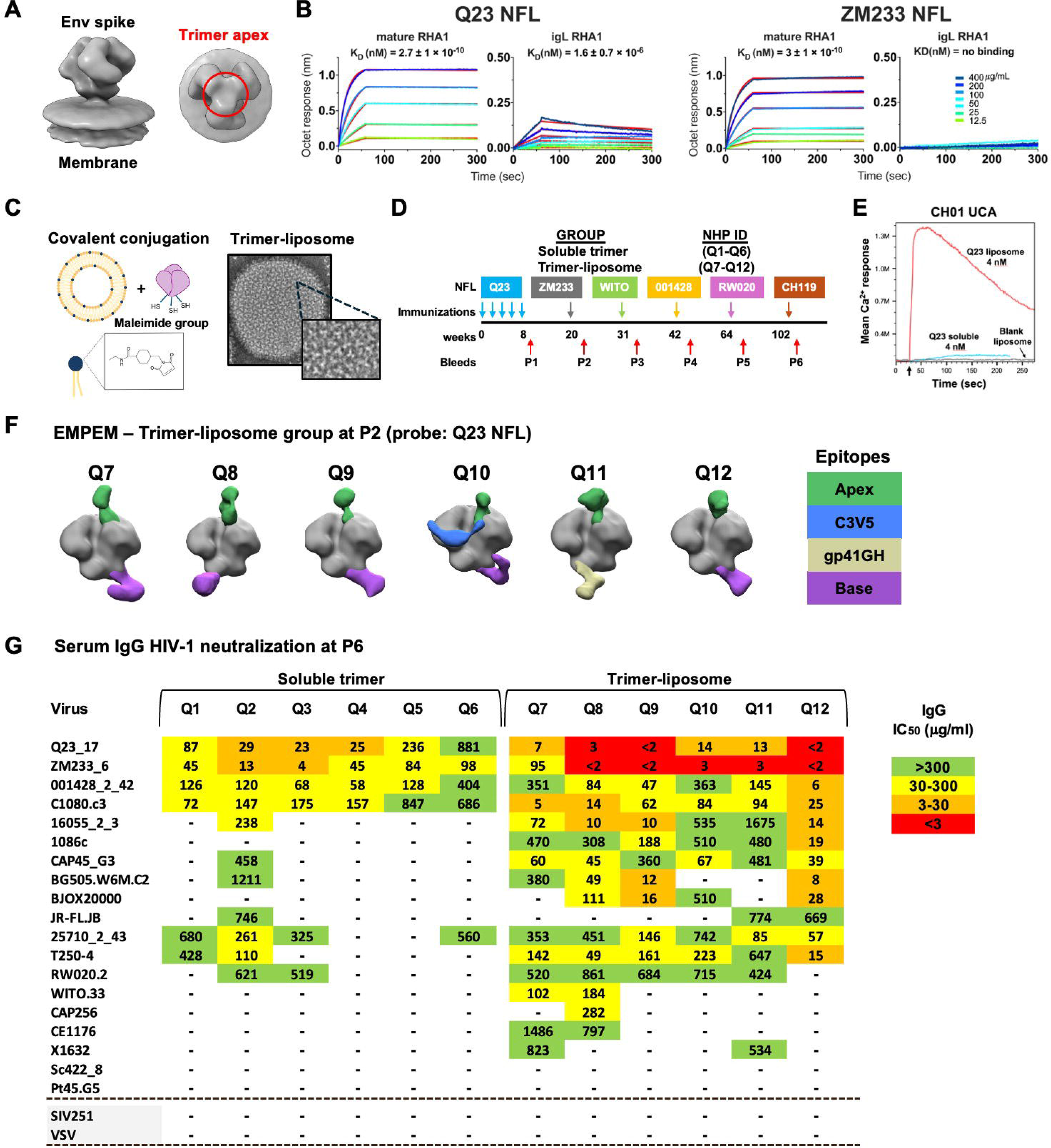
Immunization with NFL trimer-liposomes elicits tier 2 cross-neutralizing antibodies in all 6 NHPs. (A) Surface representation of HIV-1 Env trimer on the membrane (left) and trimer apex (EMD-5273) (right). (B) BLI binding analysis of Q23 and ZM233 NFL trimers to mature RHA1 and inferred germ-line reverted mAbs (igL RHA1). (C) Pictorial representation of liposomes conjugation to trimers (left). Negative stain EM image of trimer-liposome (right). (D) Immunization regimen and NFL trimers used as immunogens. (E) Calcium flux binding response of CH01 UCA to Q23 trimer-liposome or soluble trimer. (F) EMPEM analysis of Q7-Q12 NHPs after immunization with Q23 and ZM233 NFL trimers (P2 bleed). (G) Neutralizing titers against a panel of clinical isolates after immunization with six NFL trimers (P6 bleed). Neutralization experiments were done in duplicate. See also Figure S1.

Elicitation of apex-targeting bNAbs is of particular interest for vaccine design, as such antibodies often require lower levels of somatic hypermutation (SHM) to gain bNAb activity.^8^ Several potent apex-directed bNAbs have been isolated from PLWH.^9–15^ Generally, the HIV-induced apex bNAbs display very long heavy chain complementarity-determining region 3 (HCDR3) that are characterized by negatively charged ‘tips’ that interact with the basic HIV Env variable region 2 (V2) C-strand, such as those found in PG9/16, PGT145, VRC26, and CH01-4.^16–20^ The glycan-shrouded apex of the Env spike is recognized by the bNAb HCDR3s that penetrate the glycan shield interacting with both the surrounding glycans and the semi-conserved basic C-strand of V2.^8,19–22^ The maturation of these antibodies is driven by the co-evolution of the virus and antibody interactions that include both glycan and protein contacts.^21,23^

Studies show that infection with chimeric simian-human immunodeficiency viruses (SHIVs) elicit apex-targeting bNAbs in non-human primates (NHPs), such as the RHA1 bNAb, ^24^ as well as other apex-directed cross-neutralizing monoclonal antibodies (mAbs). ^25^ These findings validate macaques as a suitable model for evaluation of vaccines aimed to stimulate such responses. ^24^ The SHIV-induced cross-neutralizing serum activity detected in the RHA1-generating NHP was informative to identify virus trimers capable of eliciting apex-directed antibody responses (i.e., Q23.17, ZM233, potently neutralized). In addition, SHIV-induced broadly neutralizing apex-targeting bNAbs can inform HIV vaccine design. However, B cell activation and general immune stimulus by a replicating virus differ substantially from that induced by recombinant protein subunit vaccines. It is less certain whether subunit vaccination can elicit comparable apex-targeting neutralizing antibody responses in animal models, an important first step towards translatable human vaccine strategies.

Here, we screened a set of native-like HIV Env trimers using the apex-targeting bNAb, RHA1 and its inferred germline (gL) reverted variant (igL RHA1) to identify a potential priming immunogen. For initial priming, we selected Q23.17 HIV strain-derived native flexibly linked (NFL) stabilized trimers based upon their avid binding to the mature RHA1 bNAb and still detectable binding to the igL RHA1 mAb (Figure 1B). ^26–28^ To enhance B cell priming, we covalently arrayed these trimers on synthetic liposomes ^29^ and compared the liposome-arrayed platform with soluble NFL trimers in rhesus macaques (Figures 1C and 1D). Following Q23 NFL priming and ‘tandem’ Env boosting with the heterologous ZM233 NFL trimers, we detected robust serum neutralization of both Q23.17 and ZM233 pseudoviruses in animals from both groups of NHPs. Apex-directed electron microscopy-based polyclonal epitope mapping (EMPEM) densities were detected in all NHPs immunized with the trimer liposomal array. Following sequential boosting with two additional stabilized NFL trimers, we detected tier-2 serum cross-neutralization in all trimer-liposome immunized NHPs. Based on these encouraging results, we isolated a panel of Env-binding mAbs and characterized their genetic, functional and structural properties. CryoEM structures of mAb:trimer complexes at high-resolution showed that mAb apex-targeting mirrors the binding mode of the infection-elicited human bNAb PG9. These findings demonstrate that high-density liposome arrays of germline-guided, native-like Env trimers effectively activate apex-targeting B cells, eliciting antibody responses that recapitulate features described in human infection-driven bNAbs.

## Results

### Identification of stabilized trimers that are well-recognized by an NHP germline-reverted apex bNAb

Roark et al. (2021) demonstrated that CH505 SHIV infection generated apex-directed polyclonal responses that potently neutralized several apex-sensitive viruses, including Q23.17, ZM233.6, and T-250-4.^24^ RHA1, an apex-targeting bNAb was isolated from one of the NHPs in that study. Cross-neutralizing activity in this animal’s serum was detected within a year following SHIV infection. Based on these findings, we generated soluble, stabilized, near-native NFL trimers from several of these strains, confirmed their conformational integrity, as described here and previously by bio-layer interferometry (BLI) and differential scanning calorimetry (DSC) (Figures S1A and S1B). ^26–28,30^ NFL trimers were tested for recognition by the NHP apex-directed RHA1 bNAb and nbNAb PGT145 (Figure S1C). We observed nanomolar binding affinity of mature RHA1 to NFL trimers derived from the Q23 and ZM233 HIV-1 strains while only the Q23 NFL trimer displayed micromolar binding affinity to the reverted igL RHA1 antibody (Figure 1B). The germline-reverted RHA1 mAb was designed by reverting both light and heavy chain residues to their inferred naïve sequences, retaining only the mature HCDR3. We next evaluated whether high-avidity, multivalent Q23 NFL trimers arrayed covalently on liposomes (Figure 1C) would enhance binding to igL RHA1 as determined by ELISA. Compared with soluble trimers, liposome-arrayed Q23 NFL trimers, captured by lectin, displayed stronger binding to the igL RHA1 and to the mature PGT145, with comparable 2G12 binding, confirming that equivalent levels of trimers were captured (Figure S1D). To assess B cell activation by the NFL trimer liposomal array, we measured calcium flux in primary murine B cells expressing the unmutated common ancestor (UCA) of the infection-derived, apex-targeting human bNAb, CH01, as well as in the murine K46 cells line expressing the mature bNAb PG16. We detected a substantial increase in intracellular Ca^2+^ flux induced by the multivalent Q23 trimer-liposomes compared to soluble trimers (Figures 1E and S1E), indicating that the multivalent array could potentially activate naïve B cells more effectively.

### Immunization with Env NFL trimer-liposomes elicits apex-directed, cross-neutralizing serum antibodies in all NHPs

Twelve NHPs were immunized with Q23 NFL trimers, administered either as soluble trimers (Q1-6) or covalently coupled to high-valency array as trimer-liposomes (Q7-12) (Figure 1D). SMNP was used as the adjuvant throughout the study.^31^ To increase engagement of rare apex-bNAb precursors in the NHP repertoire, we opted for a fractionated Q23 NFL priming, dosed over 8 weeks. Antibody responses from serum-purified IgG collected after the Q23 and ZM233 tandem immunization (P2) displayed autologous neutralization of both Q23 and ZM233 pseudovirus in four NHPs from the soluble group and six NHPs from the trimer-liposome group (Figure S1F). The serum-purified IgG was concentrated to a level matching that naturally found in the serum (approximately 10 mg/mL); neutralization is reported as 50% inhibitory concentrations (IC_50_). Eventually all NHPs in both groups developed these responses over subsequent immunizations (P3-P6). Neutralizing serum antibody responses were stronger in the trimer-liposome-immunized NHPs than in the animals inoculated with soluble trimers (Figure S1E). To determine whether the neutralizing antibody responses targeted the HIV Env apex, we used electron microscopy-based polyclonal epitope mapping (EMPEM) to derive images of serum antibodies in complex with the NFL trimer immunogens. EMPEM analysis of the serum IgG-derived Fabs in complex with the Q23 trimer revealed binding densities to the trimer apex in all NHPs from the trimer-liposome group (Figure 1F) and four NHPs from the soluble group (Figure S1G). The NHPs were subsequently immunized every 12 weeks with NFL Env trimers derived from heterologous HIV strains WITO.33, 001428-2.42, RW020.2, and CH119.10. Following immunization with the 001428 NFL trimers (P4), all IgG samples from animals immunized with trimer-liposomes neutralized a sentinel panel of heterologous tier 2 HIV-1 clinical isolates (Figure S1H). The most potent cross-neutralization activity was displayed in the serum IgG derived from NHP Q12. Based on these data, we analyzed the serum by EMPEM to define the predominant cross-binding specificity. We selected three NHPs from the trimer-liposome group (Q8, Q9, and Q12) that displayed neutralization activity against several heterologous HIV strains, including 16055-2.3 and BG505.W6M. Notably, NFLs derived from these strains were not part of the immunization regimen, and we thus analyzed the pooled sera by EMPEM against these heterologous trimers. PolyFab binding densities targeting the Env apex were observed in both 16055 and BG505 trimer-Fab complexes, suggesting that the apex-directed cross-neutralization was likely mediated by the serum IgG corresponding to these EMPEM densities (Figure S1I). The serum IgG from these NHPs also neutralized other apex-sensitive clinical isolates (Figure S1H). From these data, we conclude that our vaccine strategy, prime:boosting with four NFL trimers covalently arrayed on liposomes, elicited apex-directed, cross-neutralizing antibodies in all NHPs. Additional heterologous boosting after the 00148 NFL trimer (by RW020 and CH119 NFLs, P6) generally broadened the cross-neutralizing activity of the serum IgG, particularly for Q2 in the soluble trimer group and for the entire trimer-liposome group (Figure 1G).

### mAb isolation confirms serum IgG cross-neutralization, demonstrating neutralization breadth of clinical isolates

To define the Ab reactivities responsible for the serum IgG cross-neutralization detected in the trimer-liposome-immunized NHPs, we isolated single memory B cells from peripheral blood mononuclear cells (PBMCs) collected at time points P4, P5, and P6, using Q23 and/or BG505 NFL trimers as probes for flow cytometry-based sorting (Figure 2A). We genotyped each NHP for their immunoglobulin (IG) allele repertoire using IgDiscover and assigned all heavy (HC) and light chains (LC) to the closest germline V, D, and J alleles present in that individual animal, enabling precise allele annotation and clarification of clonal relationships.^32–34^ Guided by the serum neutralization data, we focused our B cell analysis on NHPs Q9, Q10, and Q12, from which we recovered 103, 159, and 393 paired heavy and light chains, which grouped into 70, 106, and 203 clonal lineages, respectively (Figure 2A). We analyzed the sequences of the HCDR3s and selected HC and LC pairs using IGHD3-15 and having HCDR3 loops longer than 20 amino acids (aa). The core of the macaque IGHD3-15 gene contains a negatively charged motif (see (Figure 2B) that has the capacity to interact with the semi-conserved C-strand of Env V2 (residues 164-172), a region which is often basic in overall charge. ^22^ These HCDR3 features found in RHA1 are characteristic of apex-targeting bNAbs isolated from SHIV-infected NHPs. ^24,25^ The sequence analysis yielded 16, 30, and 58 HC and LC pairs from the three animals, respectively, that met these antibody signature criteria (Figure 2A).

**Figure 2.**
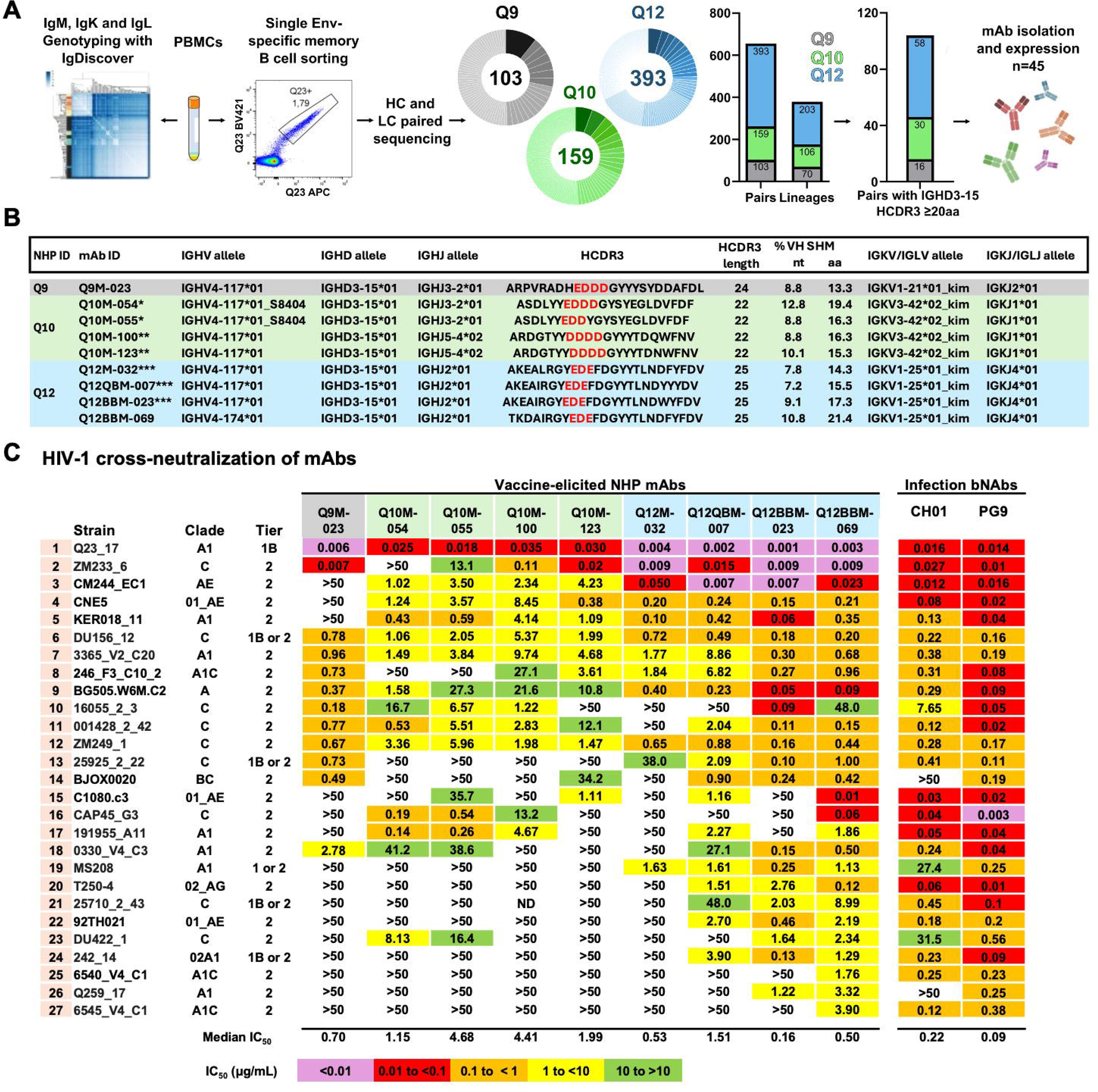
Env-specific memory B cell sorting, sequence analysis and neutralization of apex-sensitive clinical isolates by selected mAbs. (A) Schematic overview of IgG genotyping, Env-specific single memory B cell sorting, analysis of paired HC and LC sequences and expression of mAbs. (B) Genetic properties of mAbs that neutralize more than four HIV-1 strains in the initial panel (Clonally related mAbs are identified with asterisks). D3-15*01 gene central acidic residues are highlighted in red. (C) Neutralizing inhibitory concentrations IC_50_ (µg/mL) for selected mAbs against a sub-panel of 27 HIV-1 clinical isolates representing apex-sensitive viruses. The bNAbs RHA1 and PG9 are included for comparison. Neutralization experiments were done in duplicate. See also Figure S2.

Forty-five HC and LC pairs were identified to clone and express as mAbs from NHPs Q9, Q10 and Q12, and their neutralizing activity was assessed on an intial set of tier 2 HIV-1 isolates. This initial panel incorporated viruses from multiple clades that have been termed “apex-sensitive” strains due to their sensitivity of neutralization by the human infection-elicited, apex-directed bNAbs such as PG9/16, PGT145, PGDM1400, VRC26 and others^13,14,35,36^. In a few cases, these isolates are also neutralized by the germline-reverted or UCA variants of the mature apex-directed bNAbs and hence termed apex-sensitive^22,37^. Another characteristic of these isolates is the absence of the N130 glycan, which occupies space adjacent to the N160 glycan on the trimer cap, increasing antibody accessibility to the apex epitopes. In the Seaman 109-virus panel used in a previous report,^22,37,38^ 50 isolates lacked this N130 glycan. Several of the signature-defined mAbs isolated from NHPs Q9, Q10, and Q12 displayed potent autologous neutralization of the pseudoviruses Q23.17 and ZM233.6. In addition, we detected cross-neutralization of an initial panel of cross-clade HIV-1 clinical isolates (Figure S2A).

We then selected the broadest mAbs (Figure 2B) from these NHPs and tested them against a larger panel of 64 apex-sensitive viruses, including the infection-elicited apex bNAbs, CH01 and PG9, as comparative controls. The mAbs from NHP Q12 were the most broad and potent, particularly Q12BBM-069, Q12QBM-007, and Q12BBM-023, which neutralized 27, 21, and 22 isolates with median IC_50_ values of 0.5 µg/ml, 1.51 µg/ml, and 0.16 µg/ml, respectively (Figure 2C). The viruses that were not neutralized by these mAbs were further subdivided based on the presence or absence of the N130 glycan (Figure S2B). To note, some Q12 mAbs weakly neutralized the N130 glycan-containing CH505 T/F clinical isolate, but not any other isolates tested that naturally contain this apex N-glycan.

### Cryo-EM structures reveal that the vaccine-elicited mAbs target the trimer apex like the human infection-elicited bNAbs PG9 and CH03

We determined high-resolution crystal structures of four Fabs from this set, Q9M-023, Q10M-055, Q12BBM-069 and Q12QBM-007 and observed well-ordered, protruding HCDR3 β-hairpin conformation in three of the four mAbs (Figure S3A, Table S1). To define their epitope specificity at atomic detail, we analyzed these cross-mAbs in complex with NFL trimers by high-resolution cryoEM (Figure S3B, Table S2). CryoEM maps and models of the two most broadly neutralizing mAbs, Q12BBM-069 and Q12QBM-007, in complex with the BG505 NFL trimer, revealed that the vaccine-elicited mAbs bound a quaternary epitope at the trimer apex that greatly overlapped with that of the infection-induced human bNAb PG9 as shown in the BG505 SOSIP:PG9 complex structure (PDB: 7T77) (Figure 3A). The epitopes of the NHP mAbs on the trimer surface revealed a similar overall Env contact footprint, especially for the heavy chains, relative to PG9, but with a larger buried surface area (BSA) (Figure 3A). The interactions are dominated by β-hairpin HCDR3 loops that contact the glycans at position N160 in all three protomers, which also occurs with PG9 (Figure 3A). To determine the significance of these antibody-N160 glycan interactions, we generated viruses lacking the N160 glycan by genetically modifying the corresponding potential N-linked glycosylation site (PNGS). We performed a neutralization assay comparing these N160-glycan eliminated viruses to their wild type, N160 glycan-bearing counterparts, observing neutralization dependence on the presence of this glycan, consistent with the apex-directed bNAbs (Figure 3B).^13,14,35^ A similar binding pattern and N160 glycan interactions were observed in complexes of Q9M-023 and Q10M-055 with BG505 and Q23 NFL trimers, respectively (Figure S4A). The HCDR3s of all four mAbs adopted a β-hairpin motif that engaged the basic C-strand of Env V2, using both side-chain and backbone interactions, as described for the bNAbs PG9 and CH03 (Figure 4A, S4B). The acidic residues of the HCDR3 make extensive interactions with basic residues in the trimer apex (Figures 4A and S4A). In Q10M-055, a sulfated tyrosine, Y100b, at the tip of the HCDR3 interacts with the Arg169 side-chain and the Asn of N160 glycan, stabilizing its HCDR3 loop that is partially disordered in the unliganded crystal structure. This sulfated tyrosine-Env interaction occurs with the most distal Env protomeric unit increasing the antibody’s reach despite possessing a shorter HCDR3 (Figure S4A, right panel). A comparison of the cryo-EM structures of the mAbs Q12BBM-069 and Q12QBM-007 trimer complexes and the corresponding unliganded antibody crystal structures suggest that the HCDR3 β-hairpins undergo lateral shifts to avoid steric clashes to facilitate interactions with the N160 glycan and surrounding Env residues (Figure 4B, S4C). This shift varies by antibody, as measured by Cα distance between equivalent residues at the tip of the loop: 7.3 Å (Q12BBM-069), 2.8 Å (Q12QBM-007), and 2.3 Å (Q9M-023), suggesting that the elicited antibodies may have been selected in part for this flexibility, especially Q12BBM-069.

**Figure 3.**
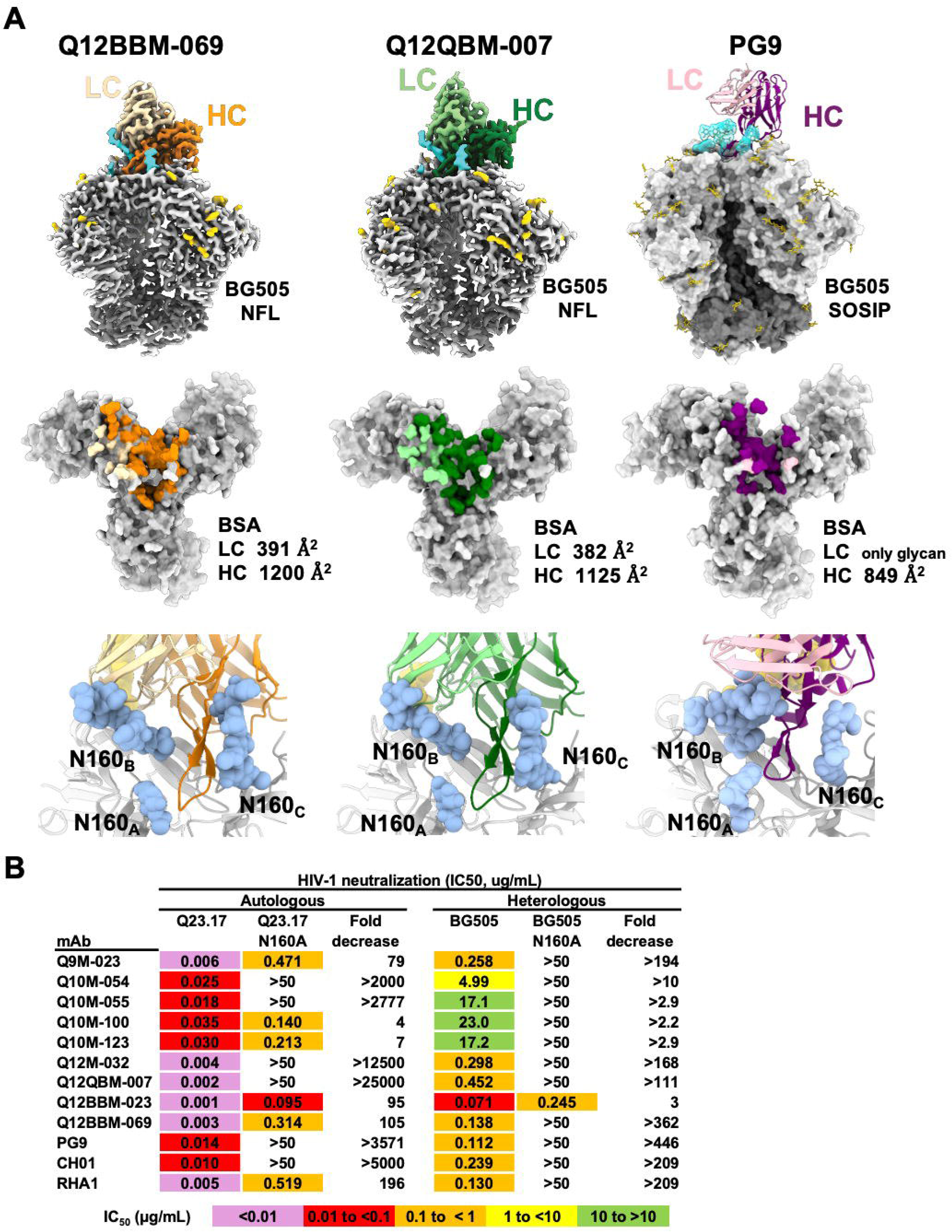
CryoEM structures of mAbs from Q12 reveal binding at the trimer-apex. (A) CryoEM maps and models of Q12BBM-069 and Q12QBM-007 Fabs in complex with BG505 NFL compared to a cryoEM model of the bNAb PG9 bound to ApexGT3.N130 BG505 SOSIP trimer (PDB 7T77) (top row). Epitope footprint of mAbs on the surface of Env trimer with buried surface area (BSA) measurements (middle row). Interaction of the HCDR3 loop of the mAbs with glycans at the N160 position (blue spheres, bottom row). (B) Neutralizing titers against Q23.17 and BG505.W6M with and without the N160 glycan. Neutralization experiments were done in duplicate. See also Figure S2.

**Figure 4.**
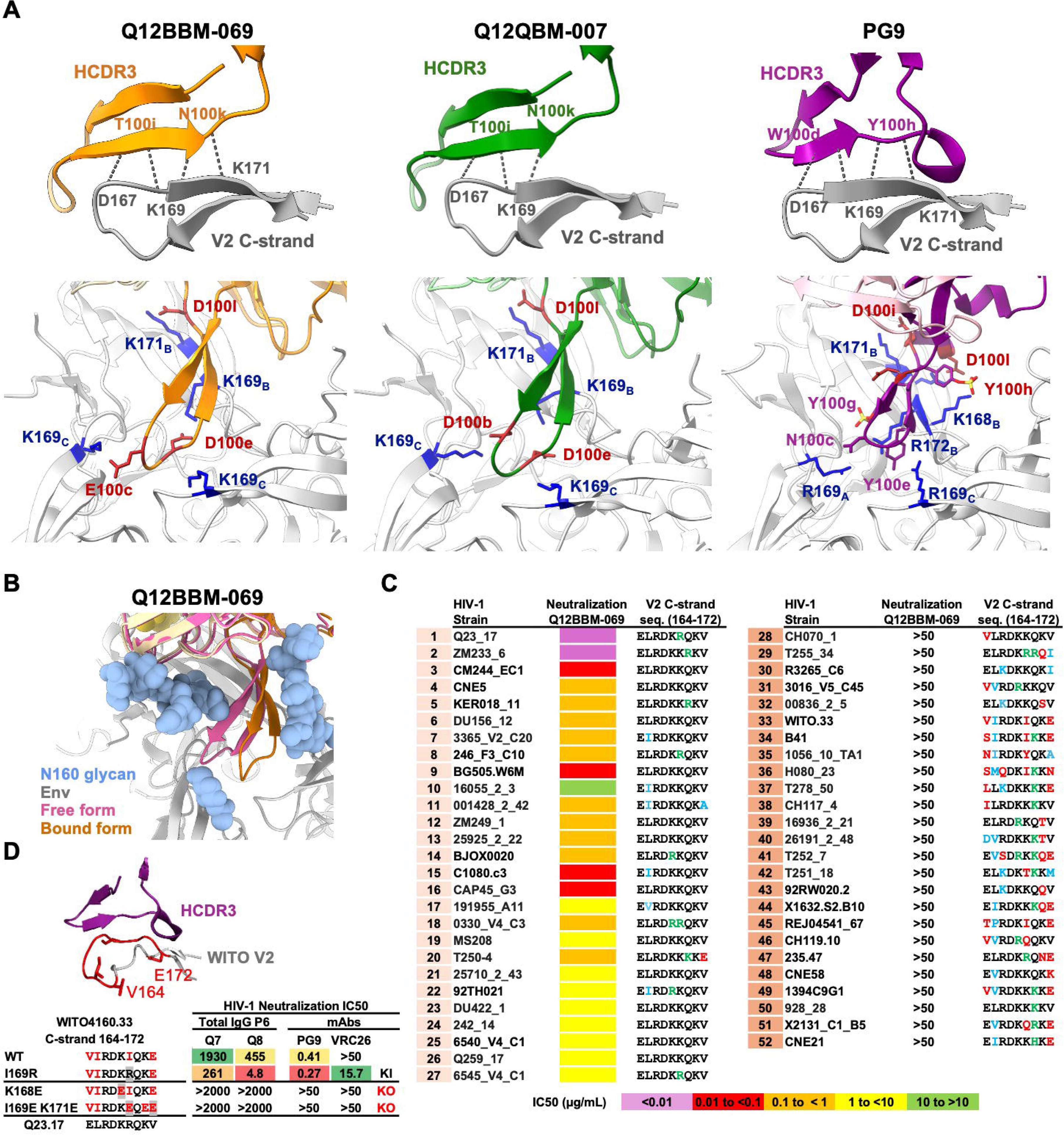
Interactions of the HCDR3 loops of NHP mAbs with the V2 C-strand at the trimer apex. (A) The β-hairpin loops of the HCDR3 of Q12BBM-069 and Q12QBM-007 interact with the C-strand of Env via hydrogen-bonding. Similar interactions are observed for the bNAb PG9 (PDB 7T77) (top row). Key interactions of the HCDR3 tip with the Env trimer-apex (bottom row). (B) Superimposition of the unbound Q12BBM-069 Fab (X-ray) with the bound form (cryoEM). (C) C-strand sequences of viruses neutralized by (left) and not neutralized (right) by Q12BBM-069 (D) Ilustration of PG9 HCDR3 interacting with the V2 C-strand of WITO Env (PDB 7T77 and PDB 9BF6) (top). Neutralizing IC_50_ titers against WITO.33 viral variants including C-strand knockout mutant viruses (bottom). Neutralization experiments were done in duplicate. See also Figure S2.

### HIV Env V2 C-strand mapping analysis provides insights into Ab cross-neutralizing specificity

To better understand the neutralization specificities of the cross-reactive mAbs isolated here, we analyzed the sequences of the HIV Env C-strand (residues 164-172) of the N130 glycan-lacking viruses tested in this study. The C-strand amino acid signature of viruses neutralized by Q12BBM-069, the most broadly neutralizing of the antibodies isolated in this study, included the following residues 164E, 165L/I, 166R, 167D, 168K/R, 169K/R, 170 Q/R/K, 171K and 172 V/A/E or “_164_ELRDKKQKV_172_”as defined by the most frequent aa at each position. This V2 sequence was highly represented in our series of NFL immunogens. The isolated mAbs make electrostatic interactions with basic amino acids (K/R) often present at four of these aa positions (166, 168, 169 and 171) (Figure 4A). Divergence from this sequence provides natural resistance to neutralization (Figure 4C). This analysis reveals that there is natural resistance in multiple isolates reminiscent of that observed by the apex bNAb VRC26, which does not neutralize clade B viruses. ^10^ Envs with C strands that differed by two or more positions as defined by the C-strand signature, like that of WITO.33, are neutralization resistant to the NHP antibodies. WITO.33 C-strand aa sequence “_164_VIRDKIQKE_172_” diverges substantially to that of the viruses neutralized by the antibodies isolated here (divergent residues in red).

While the mAbs characterized here do not neutralize WITO.33, serum IgG from NHPs Q7 and Q8 do neutralize WITO.33, a clade B virus with a divergent V2 C-strand (Figure4D). Mapping the serum IgG from Q7 and Q8 demonstrated that the observed WITO cross-neutralizing activity was directed to the V2 C-strand. Importantly, we precisely mapped this WITO.33 neutralization activities of the Q7 and Q8 serum IgG to the apex by generating WITO.33 C-strand ‘knockout’ (KO) viral variants that eliminate neutralization activity of apex bNAbs (a double mutant I169E K171E and a single mutant K168E) (Figure 4D). No neutralization activity was observed with either of these KO viruses. In contrast, a single I169R mutation in the divergent WITO.33 V2 sequence increased the neutralization sensitivity to Q7 and Q8 polyclonal IgG, highlighting the importance of the residue 169 in the apex-targeted antibody response elicited in this study. Taken together, these results indicate that neutralization of the divergent clade B WITO.33 virus is directed primarily to the C-strand and that natural resistance afforded by sequence variability in this region can be overcome. This information will guide design of improved immunization regimens based on Env trimer-liposomes. Notably, the serum IgG from NHPs Q9, Q10, and Q12 neutralized viruses 1086c and RW020.2 containing divergent C-strand sequences, indicating the presence of alternate antibody lineages in these NHPs, either directed to the apex or, potentially, targeting other epitopes.

## Discussion

In this study, we describe a vaccine strategy using near-native NFL-stabilized Env trimers covalently arrayed on liposomes that elicited apex-directed cross-neutralizing antibody responses against HIV. Our priming immunogen, Q23 NFL, was identified based on its natural and rare capacity to engage the germline-reverted apex-targeting bNAb RHA1. Sequential heterologous boosting of liposome-arrayed trimers generated cross-neutralizing antibody responses in all NHPs. Following memory B cell sorting with native-like trimers, we isolated mAbs with long, negatively charged HCDR3s (20 residues or longer), typical of apex-targeting NHP bNAbs. We demonstrate that a subset of these mAbs cross-neutralized a large panel of clinical isolates representing different HIV clades. High resolution cryo-EM structures of these antibodies in complex with selected NFL trimers confirms that they target the apex of the HIV-1 Env trimer and display structural features with remarkable similarity to the human infection-elicited bNAbs, PG9 and CH03.

These results substantially advance efforts in the HIV vaccine field to elicit broadly neutralizing responses directed to the glycan-shielded HIV-1 Env trimer apex beyond the priming of bNAb precursors alone. Typically, the so-called bNAb apex precursors bind to the original engineered germline-targeted trimers but not to more native, wild-type trimers and fail to neutralize wild-type virus. ^39^ Using germline-guided, natural apex sequence trimers, the results presented here demonstrate that we are able to not just prime apex-directed precursors, but we also educate B cells to take the next several steps by heterologous trimer boosting to drive affinity maturation of apex-directed antibodies that cross-neutralize a wide array of tier 2 cross-clade clinical isolates. The antibodies elicited here recognize wild-type Env, bind and neutralize the autologous viruses already at the post-2 time point, and then mature to develop cross-neutralizing activity following heterologous trimer boosting. We would assert this approach is an alternative to and, in this setting, may be preferable to remodeling the immunogen to become high affinity by altering the Env binding site to increase priming efficiency. Under such circumstances, achieving antibody wild-type Env recognition requires ‘boosting energy’ to evolve and in the best case, recognize the native Env trimer to achieve neutralization. In contrast, our vaccine strategy efficiently generates antibody responses capable of neutralizing autologous virus immediately following the two ‘tandem priming immunogens’.

The molecular mimicry observed between the vaccine-elicited apex cross-neutralizing mAbs presented here and the infection-driven bNAbs validates our approach. The vaccine-elicited mAbs described in this study successfully engage the N-linked glycan at position 160, an important determinant in neutralization by bNAbs PG9, PGT145, CH01, and CAP256.^22^ Extensive engagement of glycans in the HIV Env trimer by cross-neutralizing antibodies is rarely achieved by vaccination.^40^ Neutralization primarily of N130 glycan-lacking viruses may be a result of the use of trimer immunogens that also lack this N-glycan, suggesting alternative strategies moving forward. We speculate that by structure-guided immunogen modification it is possible to expand neutralization breadth by using a wider array of C-strand sequences to better evolve this process. In this regard, we detected neutralization of the clade B WITO.33 HIV strain by polyclonal serum IgG from two NHPs, suggesting that this type of antibody evolution by vaccination is possible. The capacity to elicit slightly different specificities in outbred animals shows the potential of the repertoire to elicit many such antibodies from the structural basics of the V2 apex signature with slightly different solutions and the potential to become bNAbs.

This proof-of-principle study establishes a template for further improvements with this new vaccine-focused positive control system that can be optimized for improvement by head-to-head single variable experiments in NHPs. Transitioning to humans can be accomplished similarly by using UCAs or gL-reverted bNAbs to select natural Env sequences that activate precursor B cells in the human antibody repertoire.

## Resource availability

### Lead contact

Information and requests for resources should be directed to and will be fulfilled by the lead contact, Richard T. Wyatt, email –

### Materials availability

All constructs and antibodies generated in this study are available from the lead contact without restriction.

### Data and code availability

The HC VDJ and LC VJ sequences of the Env-specific mAbs have been deposited in GenBank under the codes: For the Heavy Chains: PX281429-PX281473; for the Kappa Chains: PX281474-PX281512; and For the Lambda Chains: PX281513-PX281518. IgM, IgK and IgL repertoire data are available from ENA under the codes: Q9 IGM ERR15498913, Q9 IGK ERR15498914, Q9 IGL ERR15498915, Q10 IGM ERR15498916, Q10 IGK ERR15498917, Q10 IGL ERR15498918, Q12 IGM ERR15498919, Q12 IGK ERR15498920, Q12 IGL ERR15498921. The IgDiscover software can be found at https://gkhlab.gitlab.io/igdiscover22/. Cryo-EM maps have been deposited in the Electron Microscopy Data Bank (EMDB) under accession codes EMD-72009, EMD-72031, EMD-72033 and EMD-72035, and cryo-EM models have been deposited in the Protein Data Bank (PDB) under accession codes 9PY5, 9PYD, 9PYK and 9PYH. X-ray crystal structures of the Fabs have been deposited in the PDB under the codes 9PYN, 9PYY, 9PZ2 and 9PZ3. Representative negative stain EMPEM maps have been deposited into the Electron Microscopy Data Bank under accession codes EMD-72735, EMD-72736, EMD-72737, EMD-72738 and EMD-72739. All deposited data are publicly available. This paper does not report original code. Any additional information required to reanalyze the data reported in this paper is available from the lead contact upon request.

## Acknowledgements

The work was supported by HIVRAD grants P01 AI104722, P01 AI157299, P01 AI124337, Scripps CHAVD UM1 AI144462 and the James B. Pendleton Trust. This research was funded in part by a Distinguished Professor grant from the Swedish Research Council (agreement number 2017-00968) and the Emory National Primate Research Center Grant Nos. ORIP/ODP51OD011132 and U42PDP11023. The Emory National Primate Center is supported by the National Institutes of Health, Office of Research Infrastructure Programs/OD (P51OD011132 and U42PDP11023). We thank Theresa Fassel from Scripps Research Microscopy Core for assistance in imaging the covalent NFL trimer-liposomes and Raiees Andrabi for confirmatory gl-reverted RHA1.

## Author contributions

J.G., S.B., G.B.K.H. and R.T.W. conceived and designed the study. J.G. designed all the immunogens for this study. S.B., J.G., R.T.W. oversaw the immunization studies. S.B. designed the trimer-liposomes and purified the immunogens and trimer-liposomes. R.W. performed serum and IgG neutralization analysis. M.A., F-A.S., I.Z., X.C.D., and M.M. performed IG genotyping, B cell sorting, monoclonal antibody isolation, characterization and engineering. S.P., S.A., G.O., W-H.L., I.A.W. and A.B.W. performed or oversaw the structural analysis. D.C., B.S.H., D.L., V.R.L., D.C. and G.S. oversaw the NHP handling, immunizations, serum and PBMC sample preparation. D.J.I. provided the adjuvant., S.B., J.G., M.A., G.O., G.B.K.H. and R.T.W. interpreted the results and wrote the manuscript. All authors reviewed, edited and approved the manuscript.

## Methods

### Ethics statement

The animal work was approved by the Emory University Institutional Animal Care and Use Committee (IACUC) under protocol 202100136.

### Animals

Twelve adult Indian-origin rhesus macaques (*Macaca mulatta*) (RM) were housed at the Emory National Primate Research Center (ENPRC) and maintained in accordance with NIH guidelines. Animal care facilities are accredited by the U.S. Department of Agriculture (USDA) and the Association for Assessment and Accreditation of Laboratory Animal Care (AAALAC) International. Animals were treated with anesthesia (ketamine 5-10 mg/kg or telazol 3-6 mg/kg) and analgesics for procedures including intramuscular (IM) and subcutaneous (SC) immunization, and blood draws as per veterinarian recommendations and IACUC approved protocols. Rhesus macaques were male and female, an age range of 3-4 years old at the start of the study with an average weight of 4.8 kgs. Animals were grouped to divide age and weight as evenly as possible between the groups receiving either soluble or liposome-conjugated trimers. Animals were housed in pairs for the duration of the study.

### Cell lines

HEK293F cells were cultured in FreeStyle 293 Expression Medium (Thermo Fisher) supplemented with 1x Antibiotic Antimycotic Solution (Sigma-Aldrich) in a humidified incubator (125 rpm, 8% CO_2_, 37°C). TZM-bl, HEK293T cells were cultured in Dulbecco’s modified Eagle Medium (cDMEM) supplemented with 10% fetal Bovine Serum Albumin (BSA) and 1x penicillin-streptomycin-glutamine (Gibco) at 5% CO_2_ and 37°C. Expi293F cells were cultured in medium from 293F Expression System Kit in a humidified incubator (125 rpm, 8% CO_2_, 37°C).

### Design of NFL trimer immunogens and probes

The Env sequences used for the generation of the NFL trimeric proteins were derived from the following HIV-1 viral sequences obtained from the Los Alamos National Laboratory HIV sequence database: Q23.17 (AF0048885), ZM233.6 (DQ388517), WITO4160.33 (AY835451), 001428_2 (EF117266), 92RW020 (AY669706), BG505.W6M (DQ208456), 16055_2 (EF117268). These DNA sequences were codon optimized to enhance protein expression and modified as follows to make stabilized NFL trimeric Env proteins: the natural HIV-1 leader sequence was replaced by a CD5 leader sequence. The four-residue furin cleavage site _508_REKR_511_ was substituted with a ten-amino acid flexible linker comprising the sequence G_4_SG_4_S.^26^ The C-terminus of the Env sequence was truncated at residue 664 resulting in the elimination of the membrane proximal external proximal region (MPER), transmembrane domain (TM), and cytoplasmic tail (CT).^41^ An additional linker comprising of residues GGGGSHHHHHHHHGSGC was added to the C-terminus to facilitate coupling to liposomes via the terminal cysteine residue. Finally, a series of stabilizing mutations were introduced to create highly stable and homogeneous trimeric proteins. These stabilizing mutations consist of the TD (BG505 Trimer Derived residues), helix-breaking glycine and proline substitutions, unnatural disulfides and V3-loop and Fusion Peptide stabilizing mutations.^27,28,30^

### Trimer expression, purification, characterization and trimer-liposome preparation

The expression, purification protocols for HIV-1 Env NFL trimers were described in detail previously.^26,28,42,43^ NFL trimers used as immunogens were expressed transiently in 293F cells and purified over lectin column followed by negative selection using the non-neutralizing mAb F105 followed by size exclusion chromatography (SEC). The trimers were characterized for their structural conformation and antigenicity by SEC, Biolayer interferometry (BLI) and Differential Scanning Calorimetry (DSC). Briefly, the BLI analysis was carried out on an Octet Red instrument (Sartorius) with IgGs immobilized on anti-human IgG Fc capture sensors (Sartorius). The Env trimers were assessed as free analytes in solution (PBS pH 7.4) at a final concentration of 250 nM. Association and dissociation were measured for 60 s respectively. The data were analyzed using ForteBio software version 11.1 and kinetic parameters were obtained using a global 1:1 fit model. The thermal transition temperature (T_m_) of the NFL trimers was determined by DSC using a MicroCal VP-Capillary DSC instrument (Malvern Panalytics). The trimer samples were dialyzed in PBS, pH 7.4, and the 400 μl of the trimer sample at concentration of 0.25 mg/mL was loaded into the instrument. The dialysis buffer was used as the reference solution. The DSC experiments were performed at a scanning rate of 1 K/min under 3.0 atmospheres of pressure. The data were analyzed after buffer correction, normalization, and baseline subtraction using MicroCal VP-Capillary DSC analysis software provided by the manufacturer (Origin 7 SR4 v 7.0522). Liposome-conjugated trimers were used as immunogens in NHPs Q7-Q12. The conjugation protocol for generating trimer-liposomes was described in detail previously.^29^ Liposomes were comprised of DSPC (1,2-distearoyl-sn-glycero-3-phosphocholine), cholesterol and 1,2-dipalmitoyl-*sn*-glycero-3-phosphoethanolamine-N-[4-(p-maleimidomethyl) cyclohexane-carboxamide].^29^ Folllowing chemical conjugation with the NFL trimers, the trimer-liposomes were further purified over a Superdex 200 Increase 10/300 GL column in PBS pH 7.4 buffer to remove unconjugated trimers from the trimer-liposomes. The conjugation of trimer to liposomes was confirmed by imaging the trimer-liposomes by a negative stain electron microscope (Scripps Research, EM core).

### Calcium Flux

Spleen cells from the CH01ucaDKI mice^44^ and the K46 B cell line expressing PG16 ^45^ were used for the calcium assay. CH01uca cells were suspended at 4 million cells/ml in Advanced DMEM, labeled with 1.5 uM Calbryte^TM^ 520 AM (ATT Bioquest, #20651) with Pluronic^TM^ F-127 (Invitrogen, #P3000MP) for 30 min at 37°C. CH01uca cells were washed with Advanced DMEM and stained with TruStain FcX PLUS (BioLegend, #156604) and Alexa Fluor 647 anti-mouse B220 (BioLegend, #103226) for 10 min at room temperature. Following staining the cells were washed with 2 mM CaCl2 HBSS. After washing, cells were incubated at room temperature for 30 minutes. Two million cells were aliquoted for flow cytometric analysis. Cells were stimulated with NFL trimers or trimer-liposomes, and calcium signals were detected for 240 seconds measuring fluorescence at Ex/Em = 516/533 (B2 peak channel) on a Cytek Aurora spectral flow cytometer (Cytek Biosciences). Analysis was performed using FlowJo (Becton Dickinson). For the PG16 B cell line assay, surface expression of the BCR was induced by adding 1 μg/ml of doxycycline (Thermo Scientific, # J6380506) one day prior. The surface PG16 expression was confirmed by binding of FITC anti-human light chain lambda antibody (BioLegend, #316606) with a Cytek Aurora spectral flow cytometer (Cytek Biosciences). Further confirmation of PG16 surface expression was done by using biotinylated Q23 NFL trimer conjugated to streptavidin-Alexa Fluor 647 (Molecular probes, #S32357). Trimer or trimer-liposome induced calcium flux was measured in a similar way as done for the CH01uca spleen cells.

### Immunization of NHPs

Twelve NHPs split equally into two groups were inoculated at eight sites (four intramuscular and subcutaneous bi-lateral immunizations on deltoids and inner thighs) with 80 μg of Q23 NFL soluble trimer (NHPs Q1-Q6) or 80 μg of Q23 NFL trimer conjugated to trimer-liposomes (NHPs Q7-Q12) in 150 μg SMNP adjuvant. SMNP adjuvant was prepared as previously described^31^. NHPs were immunized with the same formulation (Q23 NFL) at weeks 2, 4, 6 and 8 as a part of the divided-dose regimen. The NHPs were boosted with 150 μg of ZM233 NFL trimer in 375 μg SMNP adjuvant at week 20. Subsequent immunizations with heterologous trimers were performed at 31, 42, and 64 weeks with 100 μg of trimer in 375 μg SMNP adjuvant. Pre-bleeds were collected prior to immunizations and test bleeds were collected on the day of immunizations and 14 days after each immunization. Blood was collected in Na Citrate CPT tubes (BD Biosciences) for peripheral blood mononuclear cells (PBMCs) and plasma isolation. Cells were cryopreserved in FBS with 10% DMSO (Gemini Bio, Fisher Bioreagents). Serum was collected via serum clot tubes (BD Biosciences).

### Neutralization

Serum IgG was affinity purified with protein A to approximately physiological levels (10mg/mL) as described previously.^30^ Replication-incompetent HIV-1 Env pseudoviruses were produced by co-transfecting HEK293T cells with 15 µg of Env-deficient backbone plasmid (pSG3Δ*env*) and 5 µg HIV-1 Env plasmid in a ratio of 1:3 (total DNA: Fugene6 transfection reagent). Pseudoviruses were harvested 72 hours after transfection and incubation at 37°C via centrifugation of cell culture supernatants at 3000g for 10 minutes and stored at −80°C. Inhibition of entry of HIV-1 Env pseudo-typed viruses into standard TZM-bl cells was used to determine the neutralization capacity of sera, purified total serum IgG or monoclonal antibodies (mAbs) as previously described in a half-well plate format.^46^ 50 µL TZM-bl cell suspension (110,000 cells/mL) in cDMEM was added to each well of white, flat-bottomed tissue culture treated plates 24 hrs before setting up the sample dilutions and incubated in a 37°C CO_2_ incubator. In a 96-well U-bottomed plate, sera, total serum IgG or antibodies were serially diluted in cDMEM five or six times (with 1:3 or 1:5 dilution factors) resulting in a total of six to seven dilution points. In a separate 96-well U-bottomed plate the sample dilutions were mixed with pre-warmed pseudovirus at a ratio of 1 to 5 (6 µL of sample dilution and 24 µL of pseudovirus), resulting in a maximal final serum dilution of 1:10, total serum IgG concentration of 2 mg/mL or antibody concentration of 50 µg/mL. The mixtures were then incubated at 37°C for one hour. cDMEM was then aspirated from the tissue culture plates containing TZM-bl cells, and 25 µL of the sample-pseudovirus mixture was pipetted directly onto the TZM-bl cells and placed in the 37°C incubator for 24 hrs. After 24 hrs 75 µL cDMEM was added to all wells of the plates and left at 37°C for another 24 hrs. Finally, the cDMEM/sample mixes were aspirated from the plates, and the TZM-bl cells were lysed for 20 minutes on an orbital shaker at 400-500 rpm. Inhibition of entry was determined using the Promega Luciferase system, with luminescence detected by a Biotek NEO2M plate reader. The resulting Luciferase signals were measured in relative light units (RLUs). Sera or antibody dilution that resulted in 50% reduction (ID_50_ values for serum and IC_50_ values for purified IgG or mAbs) in RLUs was determined by fitting the neutralization dose-response curves by non-linear regression using a 5-parameter hill slope equation. Data were analyzed using GraphPad Prism software (version 10.2.1).

### Individualized immunoglobulin genotyping

To determine the germline V, D and J allele content of the studied rhesus macaque, full-length HC VDJ and LC VJ amplicons were generated as described^33^ and analyzed by IgDiscover (https://gkhlab.gitlab.io/igdiscover22/). In brief, total RNA was extracted from approximately 5-10 million PBMCs using the Qiagen RNeasy mini kit. 300ng RNA was subjected to reverse transcription with gene-specific primers for the IgM, IgK and IgL constant regions using Sensiscript RT Kit (Qiagen), the cDNA was purified with the Qiagen MinElute PCR Purification Kit and was eluted in 20 µl of elution buffer. 3 µl cDNA were subjected to the library PCR with 25x cycles using multiplex forward VH, VK and VL primer sets and gene-specific primers for the IgM, IgK and IgL constant regions as previously described.^33^ The library was gel-purified with Qiagen MinElute Gel Extraction Kit and indexed with 10x PCR cycles following the Illumina MiSeq 2 x 300 bp kit instructions. Indexed library PCR products were purified with MiniElute PCR Purification Kit, followed by AMPure XP magnetic bead (Beckman Coulter) purification prior to sequencing with the Illumina MiSeq 2 x 300 bp kit. The libraries were analyzed using IgDiscover^34^ and Corecount^32^ to generate HC V, D and J, and LC V and J genotypes. Input databases for the IgDiscover analysis were KIMBD (http://kimdb.gkhlab.se/) for the HC alleles and IMGT for the LC alleles.

### Env-specific memory B cells sorting by flow cytometry

To produce Env trimer probes for B cell sorting, 10 µg of biotinylated Q23 and BG505 NFL trimers were conjugated to SA-APC (Invitrogen) or SA-BV421 (BioLegend) in five sequential steps, each incubation proceeded for 20 min at 4°C. Frozen single cell suspension from blood mononuclear cells (PBMCs) from animals Q9, Q10, and Q12 were thawed at 37°C, washed twice in pre-warmed RPMI 1640 media (HyClone) supplemented with 10% FBS (HyClone) and Penicillin/Streptomycin (100 IU/ 100 µg/ml) (Gibco). The cells were washed with PBS (Sigma) and counted using trypan blue exclusion of dead cells by a Countess II cell counter (Thermo Fisher). Cells were suspended in PBS and incubated for 30 minutes at 4°C with Live/Dead Fixable Aqua Dead Cell Stain Kit (Life Technologies) according to the manufacturer’s instructions. Cells were washed with FACS buffer (PBS + 1% FBS) and surface stained with the following antibodies: CD3 FITC (clone SP34-2), CD14 FITC (clone M5E2), CD20 BV421 or PerCP-Cy5.5 (clone 2H7), CD27 PE-Cy7 (clone M-T271), IgG PE-CF594 (clone G18-145) (all from BD Biosciences). Staining was performed for 30 minutes at 4°C. After washing with FACS buffer, cells were subsequently stained with the fluorescently conjugated NFL trimers. Live CD3-CD14-CD20+CD27+IgG+ENV+ single cells were sorted on a four-laser FACSAria Fusion cell sorter (Becton Dickinson) into 96-well PCR plates (Eppendorf) containing 4 µl/well of ice-cold cell lysis buffer (0.5x PBS, 10 mM DTT and 2 U/µl RNAsin (all from Thermo Fisher)). After sorting the 96-well plates were centrifuged, sealed, and immediately frozen on dry ice and stored at −80°C until use.

### Single B cell RT-PCR and mAb cloning

For cDNA synthesis the 96-well plates, containing single B cells, were thawed on ice. The reverse transcription was performed by SuperScript IV reverse transcriptase (Thermo Fisher) using random hexamers, oligodT, dNTPs (Invitrogen), Igepal CA-630 (Sigma), and RNAsin (Thermo Fisher). IgG HC and LC V(D)J sequences were amplified separately in 20 μl nested PCR reactions using 4 μl of cDNA for HC and 3 μl for LC in the 1st round PCR and 1 μl PCR product in the 2nd round PCR using KAPA HiFi HotStart ReadyMix 2x (Roche). PCR products from positive wells were purified, Sanger sequenced (Genewiz) and analyzed. HC and LC V(D)J sequences were cloned into expression vectors containing the human IgG1, Igκ1, or Igλ2 constant regions. ^47^, the sequences, engineered with overhangs complementary to the linearized vector ends, were assembled using the Gibson Assembly Master Mix (New England Biolabs). The reaction mixture, consisting of 50 ng of vector and 30 ng of insert in 20 μl reaction mix, was incubated at 50°C for one hour. After incubation, 1 µl of the diluted (1:3) reaction mix was transformed into XL10-Gold ultracompetent cells (Agilent Technologies) by heat shock at 42°C for 30 seconds. Screening of transformed colonies was assessed by PCR, positive clones were expanded, and plasmids were isolated with Plasmid Plus Midi Kit (Qiagen). The correct sequences were confirmed through Sanger sequencing (Genewiz).

### Monoclonal antibody expression and purification

Monoclonal antibodies (mAbs) were expressed by co-transfecting equal amounts of each HC and LC plasmids (18 µg each) into 30 ml FreeStyle 293-F cells (Thermo Fisher) (HEK293-F) cultured in FreeStyle 293 Expression Medium (Thermo Fisher) supplemented with 1x Antibiotic/Antimycotic solution (Sigma) at a density of 1.2 million cells/ml at >95% viability. Cultures were maintained in a humidified shaking incubator (125 rpm, 8% CO_2_, 37°C). Transfections were carried out with FreeStyle Max reagent (Invitrogen) in Opti-MEM medium (Gibco). Seven days after transfection, the mAbs were purified from the supernatant using Protein G Sepharose columns (Cytiva). 4 μg of each purified antibody was analyzed by SDS-PAGE under reducing conditions using NuPAGE 4-12% Bis-Tris polyacrylamide gels (Invitrogen).

### mAb binding to Env by ELISA

The mAbs were tested for binding to NFL Env trimers. Pierce™ NeutrAvidin™ bovine serum albumin (BSA)-blocked 96-well plates (Thermo Scientific) were coated with 5 µg/ml of NFL trimer, 50 µl/well in wash buffer (25 mM Tris, 150 mM NaCl, pH 7.2; 0.1% BSA, 0.05% Tween-20) for 2 hours. Antibodies, starting at a concentration of 10 µg/ml, were added in 25-fold serial dilutions and incubated for 1 hour at room temperature. The plates were incubated for 45 minutes at room temperature with an HRP-conjugated goat anti-human IgG (H+L) antibody (Jackson ImmunoResearch) diluted 1:10,000 in wash buffer. Each incubation step was followed by six washes with wash buffer. Binding of the mAbs was detected with TMB substrate (Life Technologies) and the reactions were stopped with 1 M sulfuric acid. Optical density (OD) measurements were performed at 450 nm using the Apollo microplate reader (Berthold).

### Electron-microscopy polyclonal epitope mapping (EMPEM)

Total Serum IgG was digested to polyclonal Fab using papain as per manufacturer’s instructions (Pierce Fab Preparation kit, Thermo Scientific). For each immune complex, 1 mg of polyclonal Fab was incubated with 15 µg of Env NFL trimer overnight and purified the next morning using a Cytiva Superdex 200 Increase size-exclusion column. Fractions corresponding to immune complexes were pooled and adsorbed onto 400-mesh carbon-coated copper grids (Electron Microscopy Sciences) at a concentration of ∼0.02 mg/mL for 10 s before blotting off the excess liquid. Grids were stained for 45 s using 2% (w/v) uranyl formate (Electron Microscopy Sciences) before blotting. Imaging was performed using an Thermo Fisher Scientific Glacios transmission electron microscope operating at 200 kV, equipped with Thermo Fisher Scientific Falcon 4 camera (73,000x magnification, 1.89 Å pixel size). Automated data collection was performed using EPU (Thermo Fisher Scientific) and data processing was performed using Relion 4.0^48^, following standard procedures. Representative maps have been deposited to the Electron Microscopy Data Bank.

### Purification and crystallization of unliganded Fabs

Fabs were expressed in Expi293F cells (Thermo Fisher Scientific, Cat. No.: A14527) transiently transfected with plasmid DNA encoding the Fab heavy and light chains at a 1:1 ratio. Transfections were performed using the ExpiFectamine 293 Transfection Kit in Opti-MEM medium. Six days post-transfection, culture supernatants were harvested and sterile filtered through a 0.22 µm membrane filter. Fabs were purified using the CaptureSelect CH1-XL affinity matrix (Thermo Fisher Scientific, Cat. No.: 1943462050), followed by size-exclusion chromatography (SEC) on a Superdex 200 16/600 column equilibrated with 20 mM Tris, 150 mM NaCl, pH 7.5. Fractions corresponding to Fab proteins were pooled and concentrated to 12 mg/mL for crystallization. Crystallization screening was performed using our Rigaku CrystalMation robotic system with JCSG Core Suites 1–4, and Top96 Cryo screens at 20°C. Crystals were cryoprotected, if necessary, by supplementing the reservoir solution with 15% ethylene glycol and then flash-cooled in liquid nitrogen for storage prior to data collection.

### X-ray data collection and structure determination

X-ray diffraction data were collected at beamline 17-ID-1 (AMX) of the National Synchrotron Light Source II. Crystals of Fabs Q9M-023, Q10M-055, Q12QBM-007, and Q12BBM-069 diffracted to resolutions of 2.34 Å, 1.63 Å, 1.54 Å, and 1.79 Å, respectively. The crystallization conditions were as follows: Q9M-023 Fab crystallized in 40% MPD, 0.1 M cacodylate buffer at pH 6.5, and 5% (w/v) PEG-8000; Q10M-055 Fab in 100 mM HEPES at pH 7.5, 10% (v/v) ethylene glycol, and 20% (w/v) PEG-8000; Q12QBM-007 Fab in 50% PEG-200 and 0.1 M citrate at pH 5.5; and Q12BBM-069 Fab in 1.26 M ammonium sulfate and 0.1 M cacodylate at pH 6.5. Data processing, including indexing, integration, and scaling, was performed using autoPROC.^49^ Structure determination was carried out by molecular replacement using Phaser within the Phenix software suite,^50^ with an initial model generated by AlphaFold 3.^51^ Subsequent model building and refinement were performed through iterative cycles using Coot and Phenix.refine, respectively.^52,53^ The quality of the final structures was assessed using MolProbity,^54^ and additional validation was carried out through the PDB validation server. Data collection and refinement statistics are summarized in Table S1.

### Cryo-electron microscopy

All Fabs used for structural analysis were prepared by cleaving the IgG using papain (Thermo Scientific Cat. #44985). The following four immune complexes were incubated overnight: a) 0.2 mg of BG505 NFL TD CC3+ with 0.25 mg of either Q12BBM-069, Q12QBM-007, or Q9M-023; b) 0.2 mg of Q23 NFL TD CC3+ with 0.25 mg Q10M-055. Complexes were purified the following morning using a HiLoad 16/600 Superdex 200 pg (Cytiva) gel filtration column. Fractions corresponding to each immune complex were pooled and concentrated using an Amicon 100 kDa MWCO centrifugal device to between 4-6 mg/mL. Samples were vitrified using a Vitrobot Mark IV (Thermo Fisher Scientific). The temperature was set to 4°C and humidity was maintained at 100% during the freezing process. The blotting force was set to 1 and wait time was set to 10 s. Blotting time was varied from 3 to 6 s. Detergent lauryl maltose neopentyl glycol (LMNG; Anatrace) was added to the sample to a final concentration of 0.005 mM shortly before freezing. UltrAuFoil 1.2/1.3 (Au, 300-mesh; Quantifoil Micro Tools GmbH) grids were used and glow discharged (PELCO easiGlow, Ted Pella, Inc.) for 40 s prior to sample application. 0.5 µL of detergent was mixed with 3.5 µL of samples and 3 µL of the mixture was immediately loaded onto the grid. Following blotting, the grids were plunge-frozen into liquid nitrogen-cooled liquid ethane.

Samples were loaded into a Thermo Fisher Scientific Glacios 2 TEM operating at 200 kV equipped with a Thermo Fisher Scientific Falcon 4i direct electron detector. Exposure magnification was set to 190,000x with a pixel size at the specimen plane of 0.718 Å. EPU software (Thermo Fisher Scientific) was used for automated data collection. Micrograph movie frames were motion corrected, dose weighted, and CTF correction was performed using cryoSPARC Live.^55^ cryoSPARC was used for the remainder of data processing. Particle picking was performed using blob picker initially followed by template picker. Particles were initially down-sampled by a factor of 4 and multiple rounds of 2D classification were performed followed by 3D ab initio reconstruction. The remaining particles were then re-extracted, with optional Fourier cropping to reduce memory requirements of large box sizes, resulting in image pixel sizes of 1.034 Å (Q12BBM-069 + BG505), 1.005 Å (Q12QBM-007 + BG505), 0.718 Å (Q9M-023 + BG505) or 1.034 Å (Q10M-055 + Q23). 3D non-uniform refinement was performed. For all cryo-EM maps, global resolution was estimated using half maps and a Fourier shell correlation cutoff of 0.143, and local resolution estimated using the cryoSPARC Local Resolution tool with 0.143 FSC cutoff.

Initial models were generated using AlphaFold3 and docked into the cryo-EM maps using UCSF ChimeraX.^51,56^ Manual building was performed in Coot 0.9.8 and real space refinement in Phenix.^57,58^ Final models were validated using MolProbity and EMRinger in the Phenix suite. Buried surface area calculations and interface analyses were performed using UCSF ChimeraX. Data collection, processing and model statistics are summarized in Figure S3 and Table S2.

---

**Figure S1.**
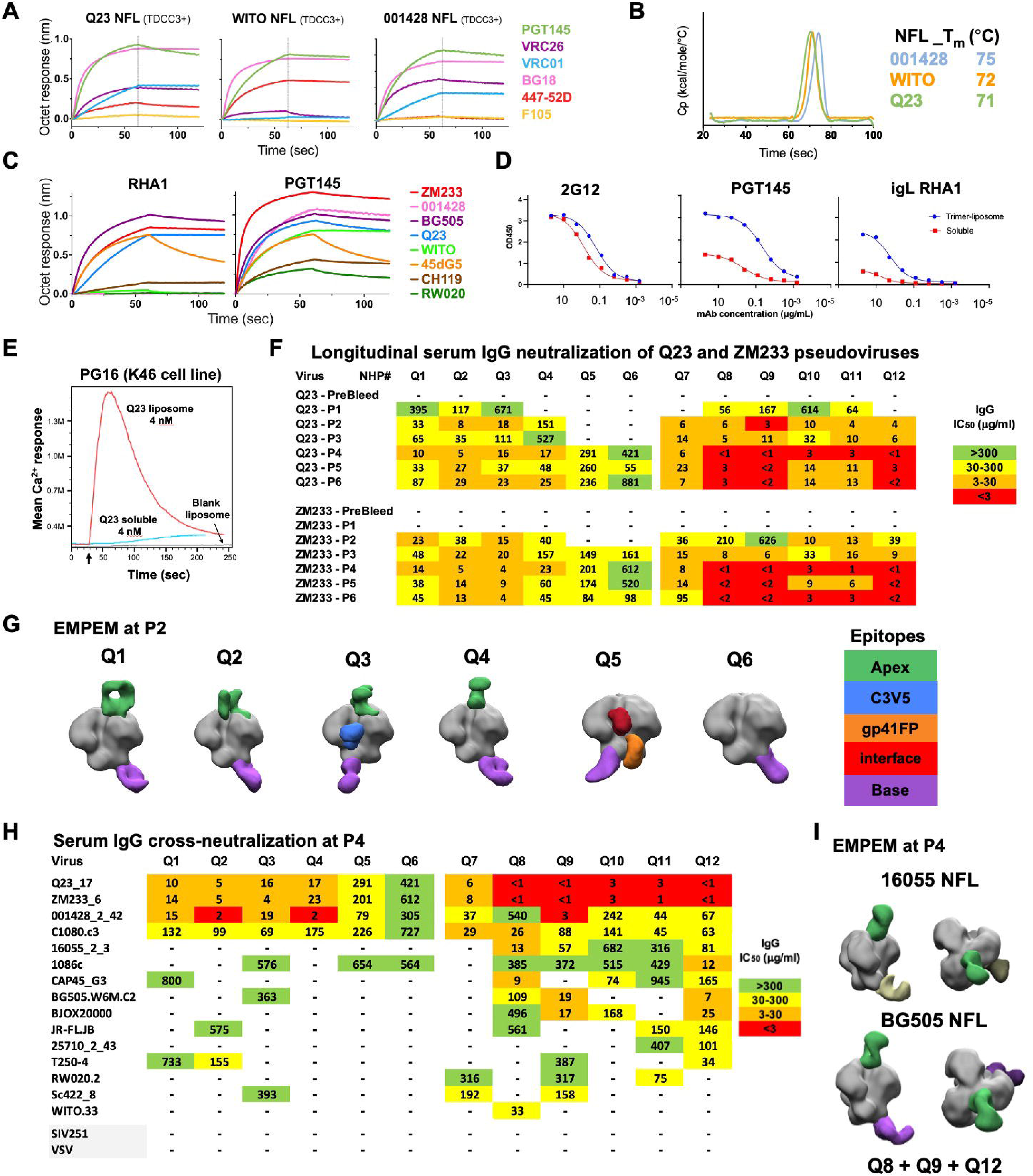
Biochemical analysis of NFL trimers, trimer-liposomes, serum and EMPEM analysis, related to Figure 1. **(A)** BLI binding of NFL trimers to representative antibodies. **(B)** DSC of NFL trimers **(C)** BLI binding of RHA1 and PGT145 to NFL trimers. **(D)** ELISA binding data of Q23 NFL trimer­ liposomes. (E) Calcium flux analysis against PG16 cells. **(F)** Longitudinal neutralization titers (IC_50_) against Q23.17 and ZM233.6 pseudoviruses. (G) EMPEM analysis of Q1-Q6 NHPs at P2. **(H)** Neutralization titers against a panel of clinical isolates at P4. (I) EMPEM analysis of pooled serum at P4 reveals cross binding density at trimer-apex (green) against two different Env trimers not from the immunogen series.

**Figure S2.**
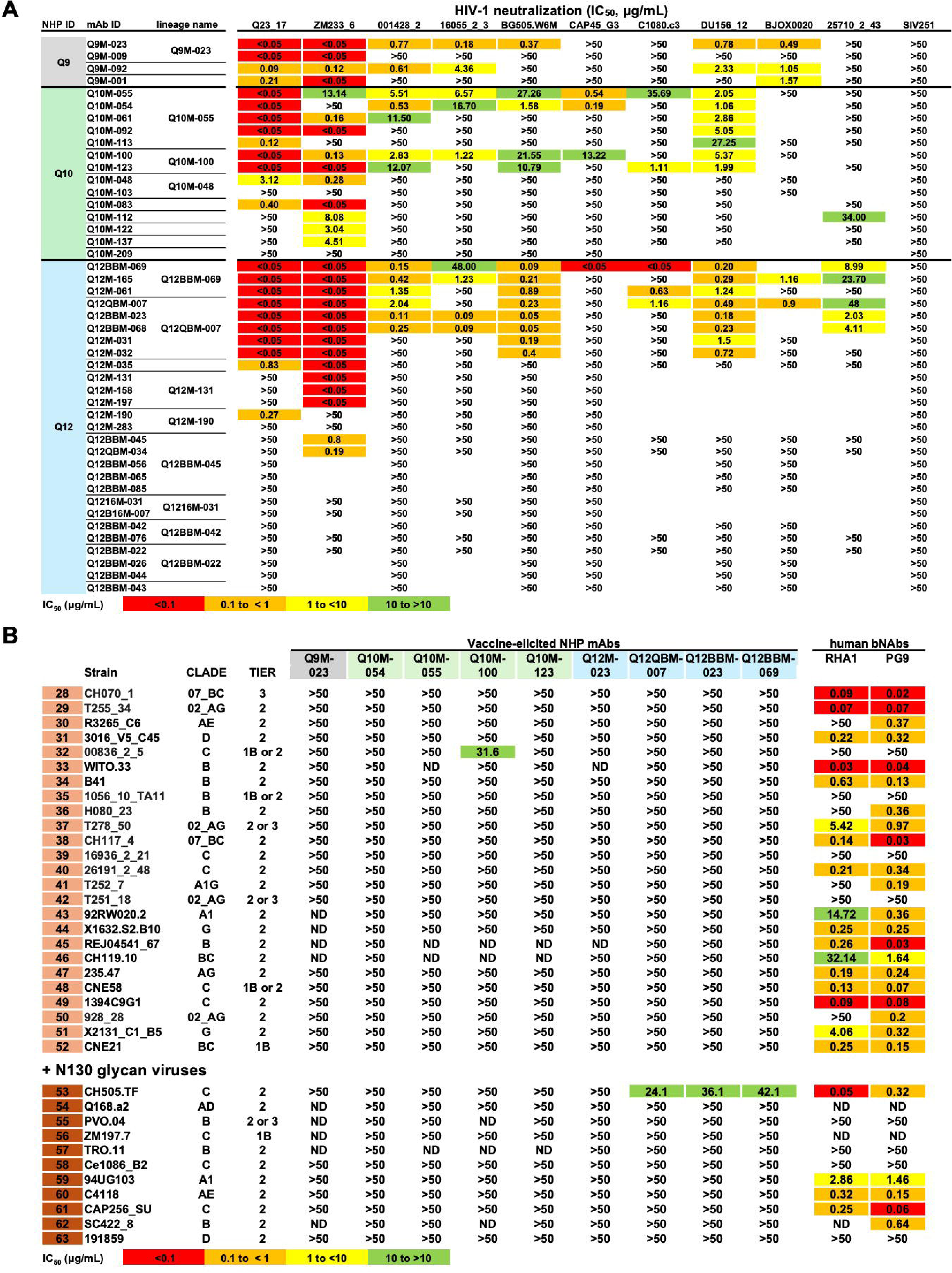
Neutralization titers of mAbs, related to Figures 1 and 2. **(A)** Initial screening of 45 mAbs cloned from Q9, Q10 and Q12 NHPs against a small panel of viruses to select mAbs with cross-neutralizing activity. **(B)** Cross-neutralizing mAbs against additional viruses lacking N130 glycan (light brown) and viruses with N130 glycan (dark brown).

**Figure S3.**
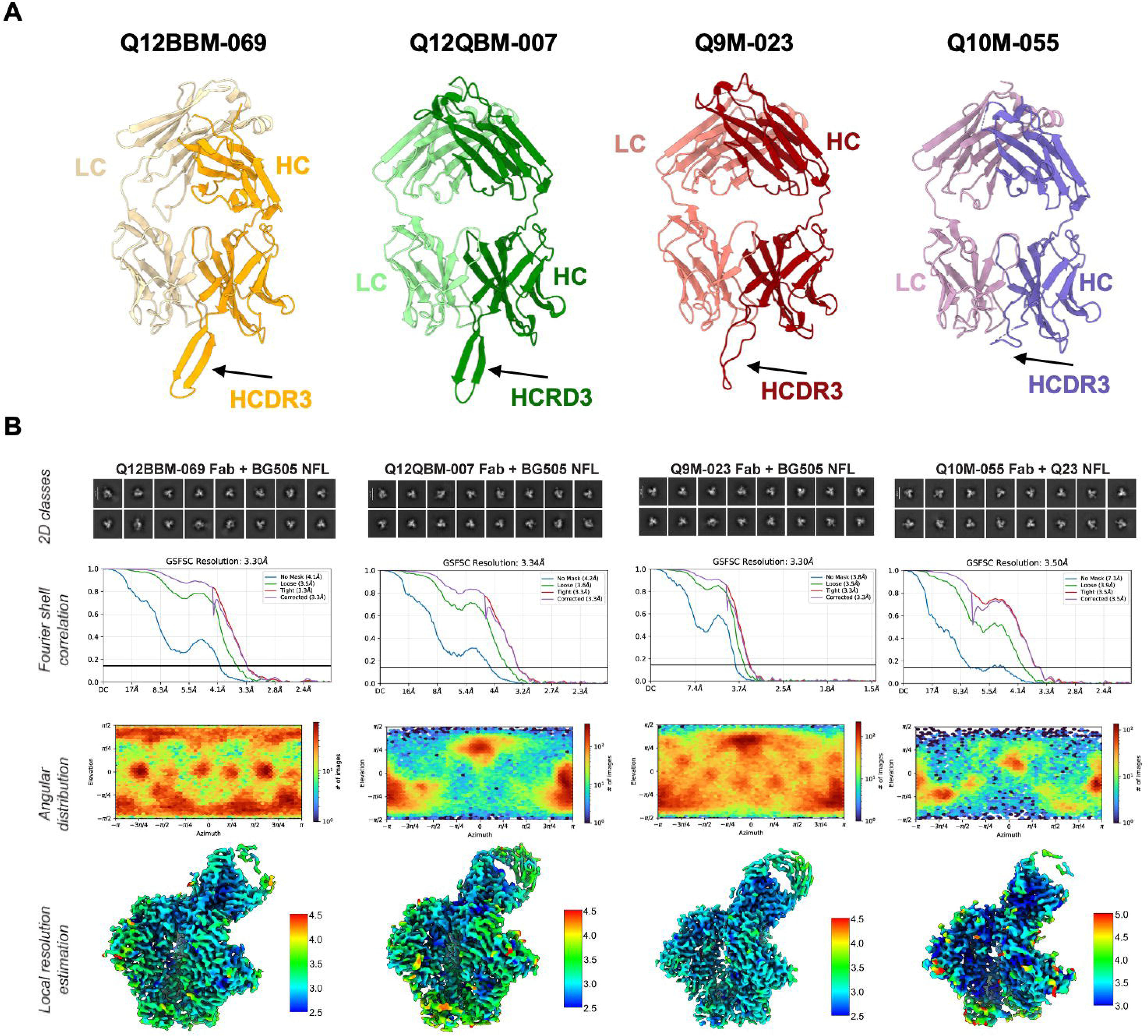
Crystal structures of Fabs and CryoEM reconstruction statistics, related to Figures 3 and 4. **(A).** Crystal structures of unliganded Fabs. HCDR3 is labeled. **(8)** Representative 2D class averages, Fourier shell correlation resolution estimate, angular distribution of observations, and local resolution estimation (in A) of the cryo-EM datasets.

**Figure S4.**
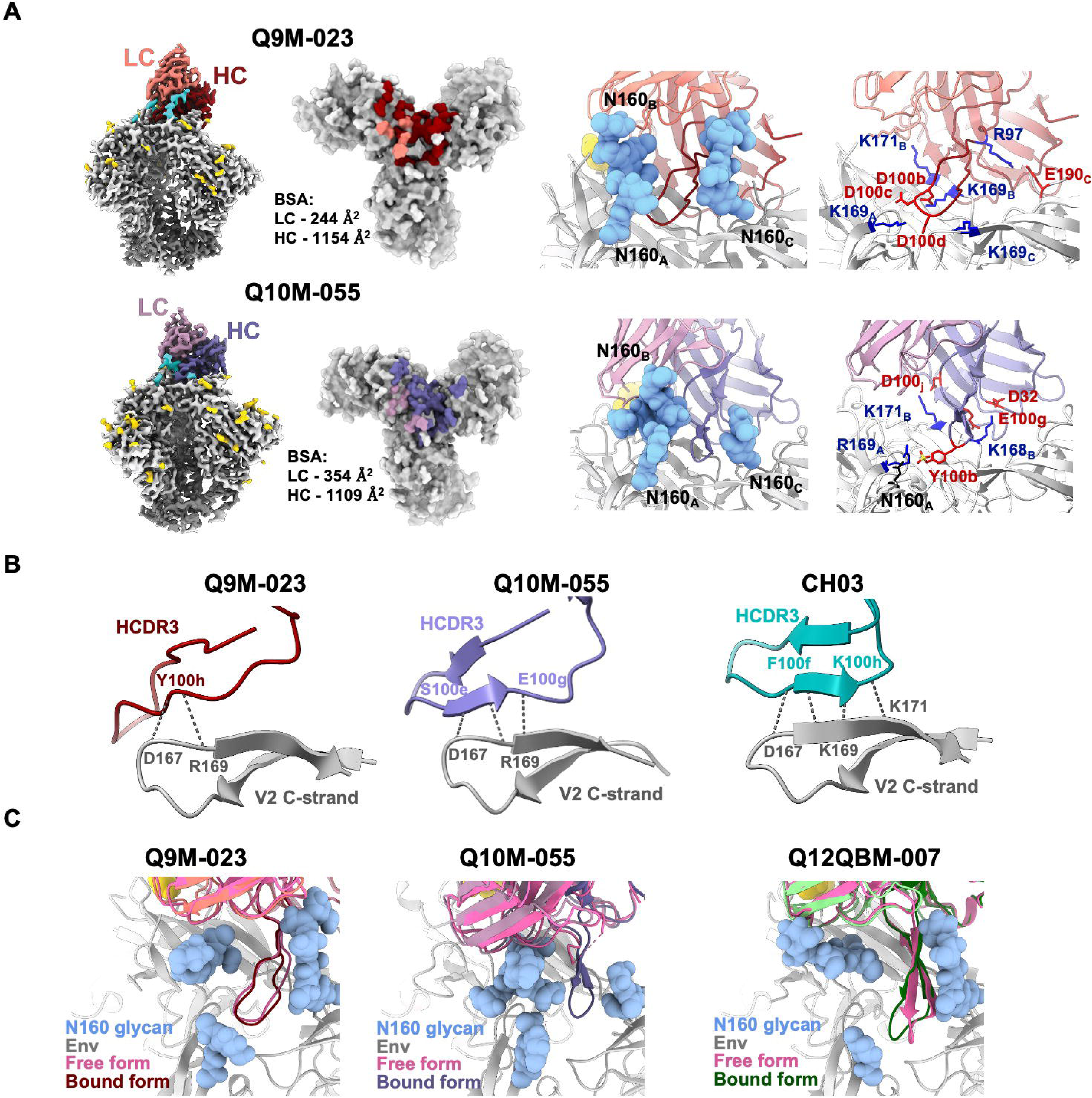
CryoEM complex structures of Q9M-023 and Q1OM-055, interaction with V2 C-strand, comparison to the unliganded HCDR3, related to Figures 3 and 4. **(A)** CryoEM maps, epitope footprint, HCDR3 interactions with N160 glycan, and key contact residues of Q9M-023 in complex with BG505 NFL (top row) and Q10M-055 with Q23 NFL. Glycans at N160 position are colored blue, all other glycans are colored yellow. **(B)** Interaction of HCDR3 with C-strand, compared to bNAb CH03. Hydrogen bond interactions are shown as dotted lines. **(C)** Superposition of HCDR3 loops from (unliganded) crystal structures to cryoEM complex structures.

