## Supplementary tables 1 and 2 for "HIV Env trimers elicit NHP apex cross-neutralizing antibodies mimicking human bNAbs"

**Table S1. X-ray data collection and refinement statistics**

| **Data Collection** | **Q9M-023** | **Q10M-055** | **Q12BQM-007** | **Q12BBM-069** |
|---|---|---|---|---|
| Beamline | NSLS-II 17-ID-1 | NSLS-II 17-ID-1 | NSLS-II 17-ID-1 | NSLS-II 17-ID-1 |
| Wavelength (Å) | 0.9197 | 0.9197 | 0.9197 | 0.9197 |
| Resolution (Å) | 33.13 - 2.34 (2.38 - 2.34) | 33.73 - 1.63 (1.66 - 1.63) | 35.14 - 1.54 (1.57 - 1.54) | 33.50 - 1.79 (1.82 - 1.79) |
| Space group | $P\ 2_1\ 2_1\ 2_1$ | $P\ 2_1\ 2_1\ 2_1$ | $P\ 2_1\ 2_1\ 2_1$ | $P\ 2_1\ 2_1\ 2_1$ |
| Unit cell a, b, c (Å) | 61.7 73.0 132.5 | 50.7 74.5 135.6 | 72.9 89.1 89.8 | 56.4 72.5 175.2 |
| α, β, γ (°) | 90 90 90 | 90 90 90 | 90 90 90 | 90 90 90 |
| Total reflections | 356,373 | 886,713 | 876,707 | 949,397 |
| Unique reflections | 26,103 | 65,019 | 84,833 | 68,718 |
| Multiplicity | 13.7 (14.1) | 13.6 (13.9) | 10.3 (5.2) | 13.8 (13.7) |
| Completeness (%) | 99.9 (99.5) | 100.0 (100.0) | 98.2 (83.3) | 99.6 (99.2) |
| Mean I/sigma(I) | 8.5 (1.2) | 15.5 (1.0) | 13.0 (1.0) | 13.9 (1.3) |
| $R_{sym}$ | 0.22 (2.5) | 0.08 (2.5) | 0.10 (1.4) | 0.12 (2.4) |
| $R_{pim}$ | 0.06 (0.71) | 0.02 (0.71) | 0.03 (0.65) | 0.03 (0.66) |
| $CC_{1/2}$ | 0.99 (0.35) | 0.99 (0.36) | 0.99 (0.32) | 0.99 (0.39) |
| **Refinement Statistics** | | | | |
| Resolution (Å) | 33.13 - 2.34 | 33.73 - 1.63 | 35.14 - 1.54 | 33.50 - 1.79 |
| Reflections total / $R_{free}$ | 26,094 / 1289 | 65,008 / 3204 | 84,729 / 4039 | 68,711 / 3407 |
| $R_{cryst}$ / $R_{free}$ | 0.23 / 0.27 | 0.22 / 0.25 | 0.19 / 0.21 | 0.21 / 0.24 |
| No. of copies in ASU | 1 | 1 | 1 | 1 |
| Number of atoms | 3367 | 3472 | 3658 | 3576 |
| macromolecules | 3347 | 3238 | 3372 | 3355 |
| solvent | 20 | 234 | 286 | 221 |
| Average B-values (Å$^2$) | 53 | 31 | 22 | 34 |
| macromolecules | 53 | 31 | 21 | 34 |
| solvent | 45 | 35 | 29 | 38 |
| Wilson B (Å$^2$) | 48 | 27 | 18 | 30 |
| **RMSD from ideal geometry** | | | | |
| Bond angle (°) | 0.5 | 1.2 | 1.3 | 1.3 |
| Bond length (Å) | 0.002 | 0.012 | 0.013 | 0.013 |
| **Ramachandran statistics (%)** | | | | |
| Favored | 97.7 | 97.4 | 98 | 97.5 |
| Allowed | 2.3 | 2.6 | 2 | 2.5 |
| **PDB Code** | **9PYN** | **9PYY** | **9PZ2** | **9PZ3** |

Statistics for the highest-resolution shell are shown in parentheses.

**Table S2.** Cryo-EM data collection, refinement and validation statistics

| | Q10M-055 Fab + Q23 NFL TD CC3+ (EMD-72009) (PDB 9PY5) | Q12QBM-007 Fab + BG505 NFL TD CC3+ (EMD-72031) (PDB 9PYD) | Q12BBM-069 Fab + BG505 NFL TD CC3+ (EMD-72035) (PDB 9PYK) | Q9M-023 Fab + BG505 NFL TD CC3+ (EMD-72033) (PDB 9PYH) |
|---|---|---|---|---|
| **Data collection and processing** | | | | |
| Microscope | TFS Glacios 2 | TFS Glacios 2 | TFS Glacios 2 | TFS Glacios 2 |
| Voltage (keV) | 200 | 200 | 200 | 200 |
| Camera | TFS Falcon 4i | TFS Falcon 4i | TFS Falcon 4i | TFS Falcon 4i |
| Collection mode | Counting | Counting | Counting | Counting |
| Magnification | 190,000x | 190,000x | 190,000x | 190,000x |
| Pixel size at detector (Å) | 0.718 | 0.718 | 0.718 | 0.718 |
| Total electron exposure (e–/Å$^2$) | 45.2 | 44.9 | 45.0 | 45.0 |
| Exposure rate (e-/pixel/sec) | 7.56 | 6.68 | 7.30 | 7.84 |
| Number of EER frames | 40 | 40 | 40 | 40 |
| Defocus range (µm) | -0.8 to -1.7 | -0.8 to -1.7 | -0.8 to -1.8 | -0.8 to -1.8 |
| Automation software | EPU | EPU | EPU | EPU |
| Micrographs collected (no.) | 7,036 | 7,003 | 7,001 | 7,021 |
| Micrographs used (no.) | 4,663 | 5,551 | 6,207 | 6,766 |
| Initial particle images (no.) | 425,148 | 947,031 | 790,420 | 645,545 |
| Final particle images (no.) | 24,541 | 81,859 | 72,419 | 131,528 |
| Map pixel size (Å) | 1.034 | 1.005 | 1.034 | 0.718 |
| Symmetry | C1 | C1 | C1 | C1 |
| Map resolution (masked/unmasked Å) | 3.5/7.1 | 3.3/4.2 | 3.3/4.1 | 3.3/3.8 |
| FSC threshold | 0.143 | 0.143 | 0.143 | 0.143 |
| Map sharpening *B* factor (Å$^2$) | -45 | -62 | -66 | -87 |
| Map resolution range (Å) | 3.0-5.0 | 2.5-4.5 | 2.5-4.5 | 2.5-4.0 |
| **Refinement** | | | | |
| Initial model used (PDB code) | AlphaFold3 | AlphaFold3, 6V0R | AlphaFold3 | AlphaFold3 |
| Refinement package | Phenix real space refine | Phenix real space refine | Phenix real space refine | Phenix real space refine |
| Model resolution (Å) | 3.8 | 3.5 | 3.5 | 3.4 |
| FSC threshold | 0.5 | 0.5 | 0.5 | 0.5 |
| EMRinger score | 2.05 | 2.16 | 2.40 | 3.13 |
| CC (mask) | 0.78 | 0.79 | 0.81 | 0.82 |
| *Model composition* | | | | |
| Non-hydrogen atoms | 16,706 | 15,794 | 15,785 | 15,625 |
| Protein residues | 1,966 | 1,896 | 1,883 | 1,857 |
| Ligands | 84 | 65 | 71 | 73 |
| *Mean B factors (Å$^2$)* | | | | |
| Protein | 101 | 66 | 49 | 64 |
| Ligand | 108 | 79 | 71 | 81 |
| *R.m.s. deviations* | | | | |
| Bond lengths (Å) | 0.005 | 0.007 | 0.006 | 0.005 |
| Bond angles (°) | 0.999 | 1.279 | 1.112 | 0.879 |
| *Validation* | | | | |
| MolProbity score | 1.12 | 1.34 | 1.25 | 1.16 |
| Clashscore | 1.82 | 2.66 | 1.44 | 1.13 |
| Poor rotamers (%) | 0.81 | 0.42 | 0.54 | 0.00 |
| *Ramachandran plot* | | | | |
| Favored (%) | 96.99 | 95.82 | 94.55 | 95.19 |
| Allowed (%) | 3.01 | 4.18 | 5.45 | 4.81 |
| Disallowed (%) | 0.00 | 0.00 | 0.00 | 0.00 |
| Cβ outliers (%) | 0.00 | 0.00 | 0.00 | 0.00 |
| CaBLAM outliers (%) | 1.48 | 2.80 | 2.18 | 3.24 |
